# The functional impact of alternative splicing in cancer

**DOI:** 10.1101/076653

**Authors:** Héctor Climente-González, Eduard Porta-Pardo, Adam Godzik, Eduardo Eyras

## Abstract

Alternative splicing changes are frequently observed in cancer and are starting to be recognized as important signatures for tumor progression and therapy. However, their functional impact and relevance to tumorigenesis remains mostly unknown. We carried out a systematic analysis to characterize the potential functional consequences of alternative splicing changes in thousands of tumor samples. This analysis revealed that a subset of alternative splicing changes affect protein domain families that are frequently mutated in tumors and potentially disrupt protein protein interactions in cancer-related pathways. Moreover, there was a negative correlation between the number of these alternative splicing changes in a sample and the number of somatic mutations in drivers. We propose that a subset of the alternative splicing changes observed in tumors may represent independent oncogenic processes that could be relevant to explain the functional transformations in cancer and some of them could potentially be considered alternative splicing drivers (AS-drivers).

## Introduction

Alternative splicing provides the potential to generate diversity at RNA and protein levels from an apparently limited number of loci in the genome (Yang et al., 2016). Besides being a critical mechanism during development, cell differentiation, and regulation of cell-type-specific functions (Norris and Calarco, 2012), alternative splicing is also involved in multiple pathologies, including cancer (Chabot and Shkreta, 2016). Many alternative splicing changes recapitulate cancer-associated phenotypes by promoting angiogenesis (Vorlova *et al*, 2011), inducing cell proliferation (Yanagisawa et al., 2008), or avoiding apoptosis (Karni et al., 2007). Alternative splicing changes may originate from somatic mutations that disrupt splicing regulatory motifs in exons and introns (Jung et al., 2015; Supek et al., 2014), as well as through mutations or expression changes in core and auxiliary splicing factors, which impact the splicing of cancer-related genes (Bechara et al., 2013; Darman et al., 2015; Madan et al., 2015; Zong et al., 2014). Alterations in alternative splicing are also emerging as relevant targets of therapy (Lee and Abdel-Wahab, 2016). For instance, lung tumors with an exon skipping in the proto-oncogene *MET* respond to *MET*-targeted therapies despite not having any other activating alteration in this gene (Frampton et al., 2015; Paik et al., 2015). Alternative splicing is also important in drug resistance. For example, a proportion of non-responders to *BRAF*-targeted therapy express a *BRAF* isoform lacking exons 4–8, which encompass the RAS binding domain (Poulikakos et al., 2011). Similarly, alternative splicing of *CD19* in relation to the aberrant activity of the splicing factor *SRSF3* impairs immunotherapy in leukemia (Sotillo et al., 2015). Thus, specific alterations in splicing induce functional impacts that provide a selective advantage to tumor cells and could represent targets of therapy.

Despite the prevalence of alternative splicing in tumors and its relation to therapy, tumor progression and metastasis (Lee and Abdel-Wahab, 2016; Lu et al., 2015; Trincado et al., 2016), its functional impacts have not been exhaustively described. Alternative splicing changes can confer radical functional changes (Wang et al., 2005), remodel the network of protein–protein interactions in a tissue-specific manner (Buljan et al., 2012; Ellis et al., 2012), and expand the protein interaction capabilities of genes (Yang et al., 2016). Here we present a systematic evaluation of the potential functional impacts of alternative splicing changes in cancer samples. We described splicing changes in terms of transcript isoforms switches per tumor sample and determined the protein features and protein–protein interactions they affected. Our analysis revealed a set of isoform switches that affect protein domains from families frequently mutated in tumors, remodel the protein interaction network of cancer drivers, and tend to occur in patients with low number of mutations in cancer drivers. Furthermore, a subset of them has driver-like properties, hence could play a role in the neoplastic process independently of or in conjunction with mutations in cancer drivers.

## Results

### Patient-specific definition of isoform switches across multiple cancer types

To determine the potential functional impacts of alternative splicing in cancer, we analyzed the expression of human transcript isoforms in 4,542 samples from 11 cancer types from TCGA (Supplemental Experimental Procedures). We described splicing changes using transcript isoforms, as they represent the endpoint of transcription and splicing, and ultimately determine the functional capacity of cells. For each gene and each patient sample we calculated the differential transcript isoform usage between the tumor and normal samples. An isoform switch was defined as a pair of transcripts, the tumor and the normal isoforms, such that the change in relative abundance in a single patient in both isoforms was higher than the observed variability across normal samples. Moreover, the involved gene must not show differential expression between tumor and normal. Additionally, we discarded switches with a significant association with stromal or immune cell content (Supplemental Experimental Procedures). The final set of switches identified and that we kept for further analysis had a mean change in relative abundance of 54% and a standard deviation of 7%.

In all patients we found a total of 8,122 different isoform switches in 6,442 genes that described consistent changes in the transcriptome of the tumor samples and that would not be observable by simply measuring gene expression changes (Figure 1A) (Table S1). These switches occurred in 4443 patients; each switch in 5 or more patients, with the majority (75%) occurring in 10 or more patients (Table S1). Using SUPPA (Alamancos et al., 2015) we calculated the relation with local alternative splicing events (Supplemental Experimental Procedures). From the 8122 switches, 5667 (69.7%) were mapped to one or more local alternative splicing events. Compared with the expected proportion of event types, we observed an enrichment of alternative 5’ss, alternative first exon and retained intron, and a depletion of alternative 3’ss, alternative last exon, mutually exclusive exons and exon-cassette (Figure S1A). Mapping the tumor isoform to either form of the event, we observed that retained intron events are predominantly retained, in agreement with previous observations (Dvinge and Bradley, 2015); whereas exon-cassette events were predominantly skipped (Figure S1B). Interestingly, 30,3% of the switches were not mapped to any event, indicating that transcripts provide a wider spectrum of RNA variation compared to local alternative splicing events.

**Figure 1.**
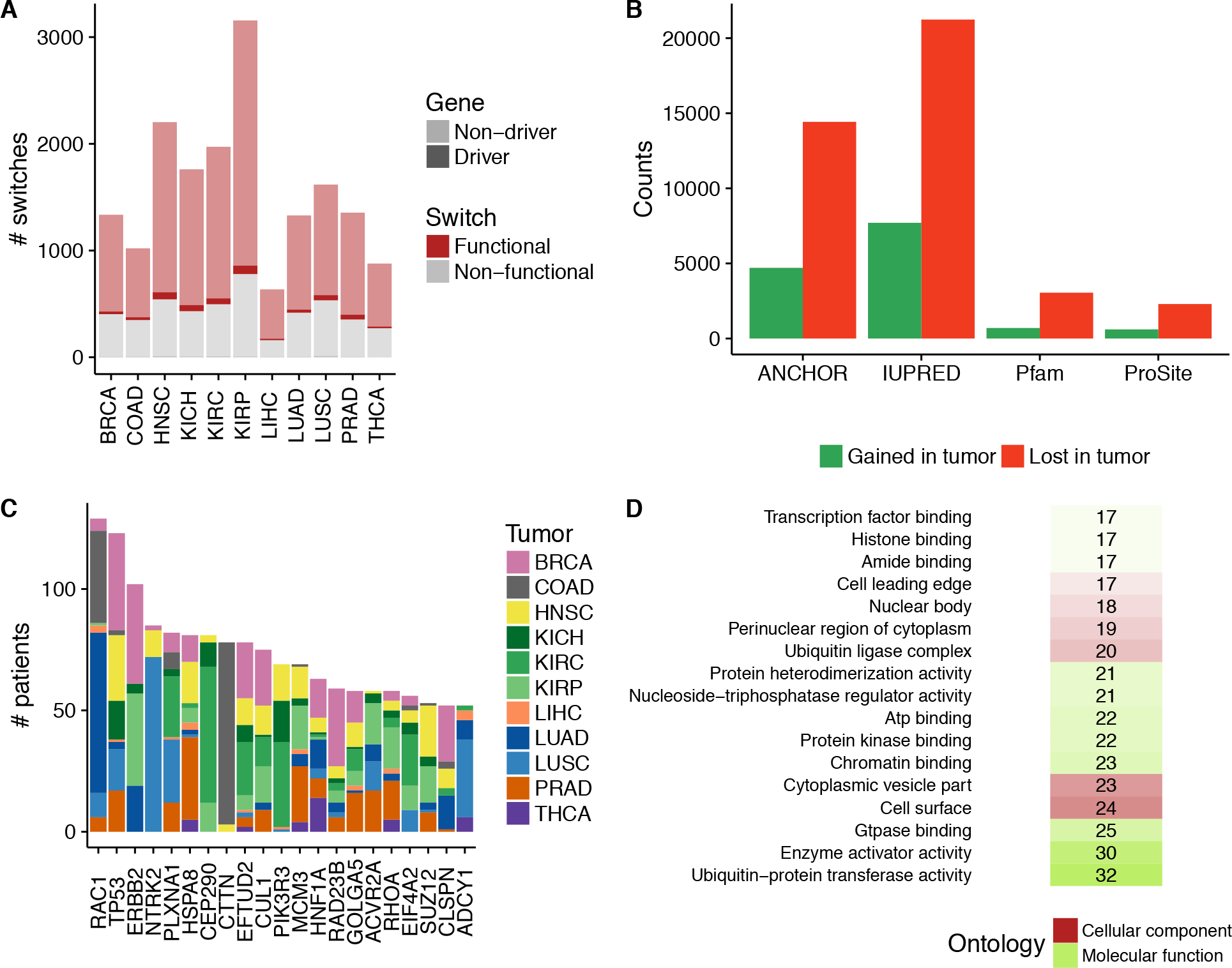
Patient-specific definition of isoform switches across multiple cancer types. **(A)** Number of isoform switches (*y* axis) calculated in each tumor type, separated according to whether the switches affected an annotated protein feature (Functional) or not (Non-functional) and whether they occur in cancer gene drivers (Driver) or not (Non-driver). **(B)** Number of different protein feature gains and losses in functional switches for each of the protein features considered, which showed significant enrichment in losses compared to random switches: Pfam (Fisher’s exact test p-value = 4.4-23, odds-ratio (OR) = 1.5), Prosite (p-value = 1.4e-08, OR = 1.3), IUPRED (p-value = 1.1e-127, OR=1.3), ANCHOR (p-value = 7.5e-139, OR=1.5). **(C)** Top 20 functional switches in cancer drivers (*x* axis) according to patient count (*y* axis). Tumor types are indicated by color: breast carcinoma (BRCA), colon adenocarcinoma (COAD), head and neck squamous cell carcinoma (HNSC), kidney chromophobe (KICH), kidney renal clear-cell carcinoma (KIRC), kidney papillary cell carcinoma (KIRP), liver hepatocellular carcinoma (LIHC), lung adenocarcinoma (LUAD), lung squamous cell carcinoma (LUSC), prostate adenocarcinoma (PRAD), and thyroid carcinoma (THCA). **(D)** Cellular component (red) and Molecular function (green) ontologies associated with protein domain families that are significantly lost in functional isoform switches (Binomial test - BH adjusted p-value < 0.05). For each functional category, we give the number of switches in which a domain family from this category is lost, which is also indicated by the color shade.

### Isoform switches in cancer are frequently associated with protein feature losses

We next studied the proteins encoded by the transcripts involved in switches. Interestingly, annotated proteins in tumor isoforms tended to be shorter than proteins in normal isoforms (Figure S1C). Moreover, while for most switches—6,937 (85,41%)—both transcript isoforms coded for protein, the rest had a significantly higher proportion of cases with only the normal isoform as protein-coding, 732 (9.01%) *vs*. 231 (2.8%) (Binomial test p-value < 2.2e-16, using 0.5 as expected frequency) (Table S1), suggesting that isoform switches in tumors are associated with the loss of protein coding capacity. To determine the potential functional impact of the isoform switches, we calculated the protein features they affected. Out of the 6,937 switches with both isoforms coding for protein, 5,047 (72.7%) involved a change in at least one of the following features: Pfam domains, Prosite patterns, general disordered regions, and disordered regions with potential to mediate protein–protein interactions (Figure S1D). Interestingly, there was a significant enrichment in protein features losses when compared with a set of 100 sets of simulated switches, controlling for isoform expression (Figure 1B). This enrichment was observed despite the fact that for simulated switches the normal protein isoform also tended to be longer than the tumor protein isoform (Figure S1E). This indicates that isoform switches in cancer are strongly associated with the loss of protein function capabilities.

We focused on the 6004 (73.9%) isoform switches that had a gain or loss in at least one protein feature, which we named *functional switches*, as they were likely to impact gene activity (Table S1). These functional switches included 729 (8.9%) and 228 (2.8%) cases for which only the normal or the tumor isoform, respectively, coded for a protein with one or more protein features. Interestingly, cancer drivers were enriched in functional switches (Fisher’s exact test p-value = 2.0e-05, OR = 1.9) (Figure S1F). Among the top switches in cancer drivers we identified one in *RAC1*, which was linked before to tumor initiation and progression (Zhou et al., 2012) and which we predicted to gain an extra Ras family domain; and one in *TP53* we predicted to change to a non-coding isoform (Figure 1C).

To characterize how functional switches affected protein function, we calculated the enrichment in gains or losses of specific domain families with respect their proportions in a reference proteome. To ensure that this was attributed to a switch and not to the co-occurrence of two domains, we requested a minimum of two switches in different genes affecting the domain. We detected 220 and 41 domain families exclusively lost or gained, respectively, and 13 that were both gained and lost, more frequently than expected by chance (Table S2). Domain families that were significantly lost included those involved in regulation of protein activity (Figure 1D), suggesting effects on protein–protein interactions. To further characterize these functional switches, we calculated the proportion of oncogenes or tumor suppressors that contained domain families enriched in gains or losses, compared with the reference proteome. From the 69 cancer drivers with domains enriched in gains, 58 (84%) corresponded to oncogenes (Fisher’s exact test p-value = 0.0066, OR = 0.4). Although tumor suppressors were not enriched in domain losses, domain families enriched in gains occurred more frequently in oncogenes than in tumor suppressors (Wilcoxon test p-value = 9e-04). These results suggest a similarity between our functional isoform switches and oncogenic mechanisms in cancer.

### Isoform switches and somatic mutations affect similar domain families

We conducted various comparisons using our switches and *cis*-occurring mutations from whole exome (WES) and whole genome (WGS) sequencing data (Supplemental Experimental Procedures). The frequencies of genes or samples with functional switches were similar to those with protein-affecting mutations (PAMs), but smaller than the frequencies for all mutations from WGS data (Figures S2A and S2B), indicating a similar prevalence of switches and PAMs but not for switches and WGS mutations. Since we calculated switches per patient, we were able to study how these distributed across patients (Supplemental Experimental Procedures). The top cases according to the co-occurrence of WGS somatic mutations with switches across patients included a switch in the cancer driver *CUX1*, although only in 7 patients (Figure S2C and S2D); whereas the top cases according to the number of patients with mutations and switches included *TP53* as well as *FAM19A5*, *DST* and *FBLN2*, which we already described as isoform switches before (Sebestyén et al., 2015) (Figures S2E and S2F). In agreement with the observed low association of mutations and switches (Figure S2G), the number of genes with PAMs and functional switches tended to be inversely correlated (Figure 2A), suggesting a complementarity between PAMs and switches affecting protein domains.

**Figure 2.**
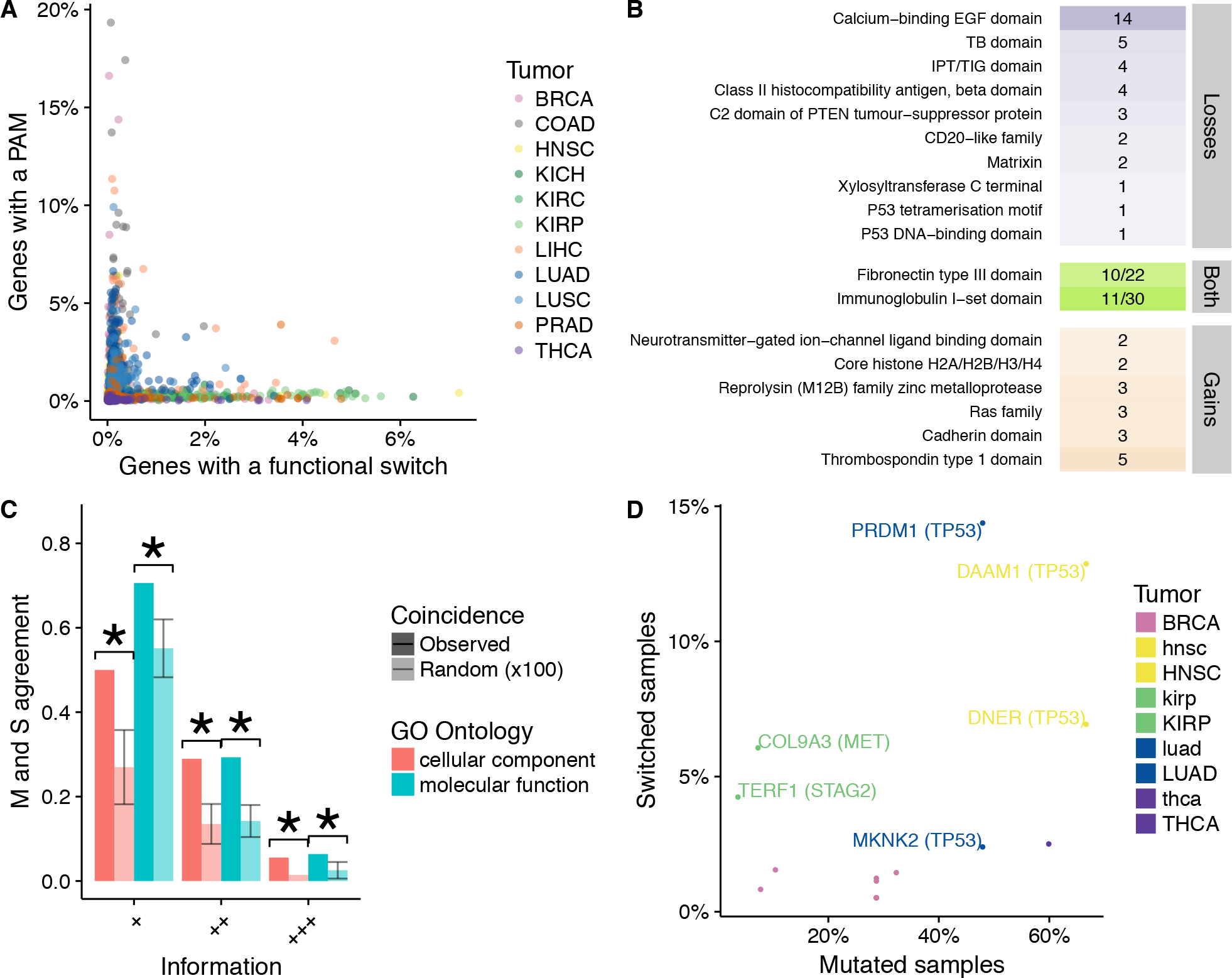
Comparison of isoform switches and somatic mutations. **(A)** For each patient sample, color-coded according to the tumor type, we indicate the proportion of all genes with protein-affecting mutations (PAMs) (*y* axis) and the proportion of genes with multiple transcript isoforms that presented a functional isoform switch in the same sample (*x* axis). (B) Domain families that were significantly lost or gained in functional isoform switches that are also significantly enriched in protein-affecting mutations in tumors. For each domain class, we indicate the number of different switches in which they occurred. We include here the loss of the P53 DNA-binding and P53 tetramerization domains, which only occurred in *TP53*. (C) Agreement between protein-affecting mutations and functional switches (*y* axis) measured in terms of the functional categories of the protein domains they affected (*x* axis), using two gene ontologies (GOs) at three different GO Slim levels, from most specific (+++) to least specific (+). Random occurrences (plotted in light color) were calculated by sampling 100 times the same number of GO terms from the reference proteome as those enriched in domain families affected by functional switches and in domains families affected by PAMs. Agreement is calculated as the percentage of the union of functional categories from both sets that are common to both. (D) Pairs formed by a cancer driver (in parentheses) and a functional switch from the same pathway and showed significant mutual exclusion (before multiple test correction) between PAMs and switches across patients in at least one tumor type-color-coded by tumor type. The *y-axis* indicates the percentage of samples where the switch occurred and *x*-axis indicates the percentage of samples where the driver was mutated in the same tumor type.

We explored this complementarity by checking if mutations and switches affected the same molecular mechanisms. First, we calculated domain families enriched in PAMs and found 76 domain families across 11 tumor types enriched in mutations (Table S2), which were more frequent in cancer drivers compared to non-drivers (Wilcoxon test p-value < 2.2e-16), in agreement with recent reports (Yang et al., 2015). Then, we compared the domain families enriched in mutations with those enriched in gains or losses through switches, we found an overlap of 15 domain families, which was higher than expected by chance given the domains affected by the 6,004 functional switches and the 5,307 domain families observed in the reference proteome (Fisher’s test p-value = 5.6e-06, OR = 4.7). From the domain families enriched in mutations, 7 showed enrichment in losses, 6 showed enrichment in gains, and 2 showed enrichment in both (Figure 2B) (Tables S2). The gains included Cadherin domains related to switches in *CHD8, CDH26, FAT1, FAT2 and FAT3*, whereas the losses included the Calcium-binding EGF domain, which is affected by various switches, including one in *NOTCH4*. A notable case was the loss of the *TP53* DNA-binding domain and the tetramerization motif. Although it occurred in a single switch, its recurrence in 123 patients highlights the relevance of *TP53* alternative splicing (Bourdon, 2007).

We questioned if the similarity was beyond the coincidence of single domain families, and could affect more generally the function associated to domains. Hence, we calculated the enriched Gene Ontology (GO) terms associated to the domains enriched in mutations and switches separately, and then calculated the overlap between both set. This overlap was compared to the overlap obtained by randomly sampling hundred times from the reference proteome the same number of GO terms found for domains in enriched switches or mutations. Notably, the observed overlap was higher than expected for each GO term and at different GO slim levels (Figure 2C), and the shared functional categories included receptor activity and protein binding. A total of 754 (12.5%) functional switches in 634 genes (47 of them in 37 cancer drivers) affected domain families that were also enriched in mutations, supporting the notion that isoform switches and mutations may impact similar functions in tumors.

If switches and mutations have similar functional impacts, we would expect a tendency toward mutual exclusion of some switches with mutations in cancer drivers. In fact, we identified 292 functional switches that were mutually exclusive with somatic PAMs in three or more cancer drivers (Fisher’s test p-value < 0.05) (Supplemental Experimental Procedures), and 16 of them showed mutual exclusion with at least one cancer gene driver from the same pathway (Table S3). These 16 switches included one in *COL9A3*, which had mutual exclusion with *MET* mutations in kidney renal papillary cell carcinoma (KIRP), and one in *PRDM1*, which showed mutual exclusion with mutations in *TP53* in lung adenocarcinoma (LUAD) (Figure 2D) as well as in *PTEN* In lung squamous cell carcinoma (LUSC) (Figure S2H) (Table S3). Despite the observed mutual exclusion, none of the cases was significant after multiple test correction, indicating that the described switches may not provide strong signatures for pan-negative tumors (Saito et al., 2015).

### Isoform switches affect protein interactions with cancer drivers

Many of the frequently lost and gained domain families in functional switches were involved in protein binding activities, indicating a potential impact on protein–protein interactions (PPIs) in cancer. To study this, we used data from five different sources to build a consensus PPI network with 8,142 nodes, each node representing a gene (Figure S3). Then, to determine the effect of switches on the PPI network, we mapped PPIs from this network to domain–domain interactions (DDIs). Domains involved in DDIs were mapped to the specific protein isoforms using their encoded protein sequence. For genes with switches, we then considered those PPIs that could be mapped to DDIs involving domains mapped on either the normal or the tumor isoforms (Figure S4). From the 8,142 genes in the PPI network, 3,243 had at least one isoform switch, and for 1,688 isoform switches (in 1,355 genes) we were able to map at least one PPI to a specific DDI with domains on either the normal or the tumor isoform. A total of 162 of these switches were located in 123 cancer drivers, with the remaining 1,526 in non-driver genes.

For each isoform switch, using the DDI information, we evaluated whether the change between the normal and tumor isoforms would affect a PPI from the network by matching the domains affected by the switch to the domains mediating the interaction, controlling for the expression of the isoforms predicted to be interaction partners. We found that 477 switches (28.3%) in 423 different genes affected domains that mediated protein interactions and thus likely impacted such interactions. Most of these interaction-altering switches (*n* = 414, 86.8%) caused the loss of the domain that mediated the interaction, while a minority (*n* = 64, 13.2%) led to a gain of the interacting domain. Only a switch in *TAF9* led to gains and losses of interactions with different partners, mediated by the loss of a TIFIID domain and a gain of an AAA domain (Table S4).

Notably, switches in driver genes tended to lose PPIs more frequently than those in non-drivers, (Figure 3A). From the 162 switches in drivers, 41 (25.3%) of them altered at least one interaction, either causing loss (33 switches) or gain (8 switches). Moreover, switches that affected domains from families enriched in mutations or that showed frequent mutual exclusion with mutational drivers also affected PPIs significantly more frequently than other functional switches (Chi-square test p-value < 2.2e-16 and p-value = 6.8e-08, respectively) (Figure S5). Looking at genes annotated as direct interactors of drivers, they tended to affect PPIs more frequently than the rest of functional switches mapped to PPIs (Figure 3B). Additionally, all functional pathways found enriched in PPI-affecting switches were related to cancer (adjusted Fisher’s exact test p-value < 0.05 and odds-ratio > 2) (Table S5), reinforcing the functional relevance of these 477 PPI-affecting isoform switches in cancer.

**Figure 3.**
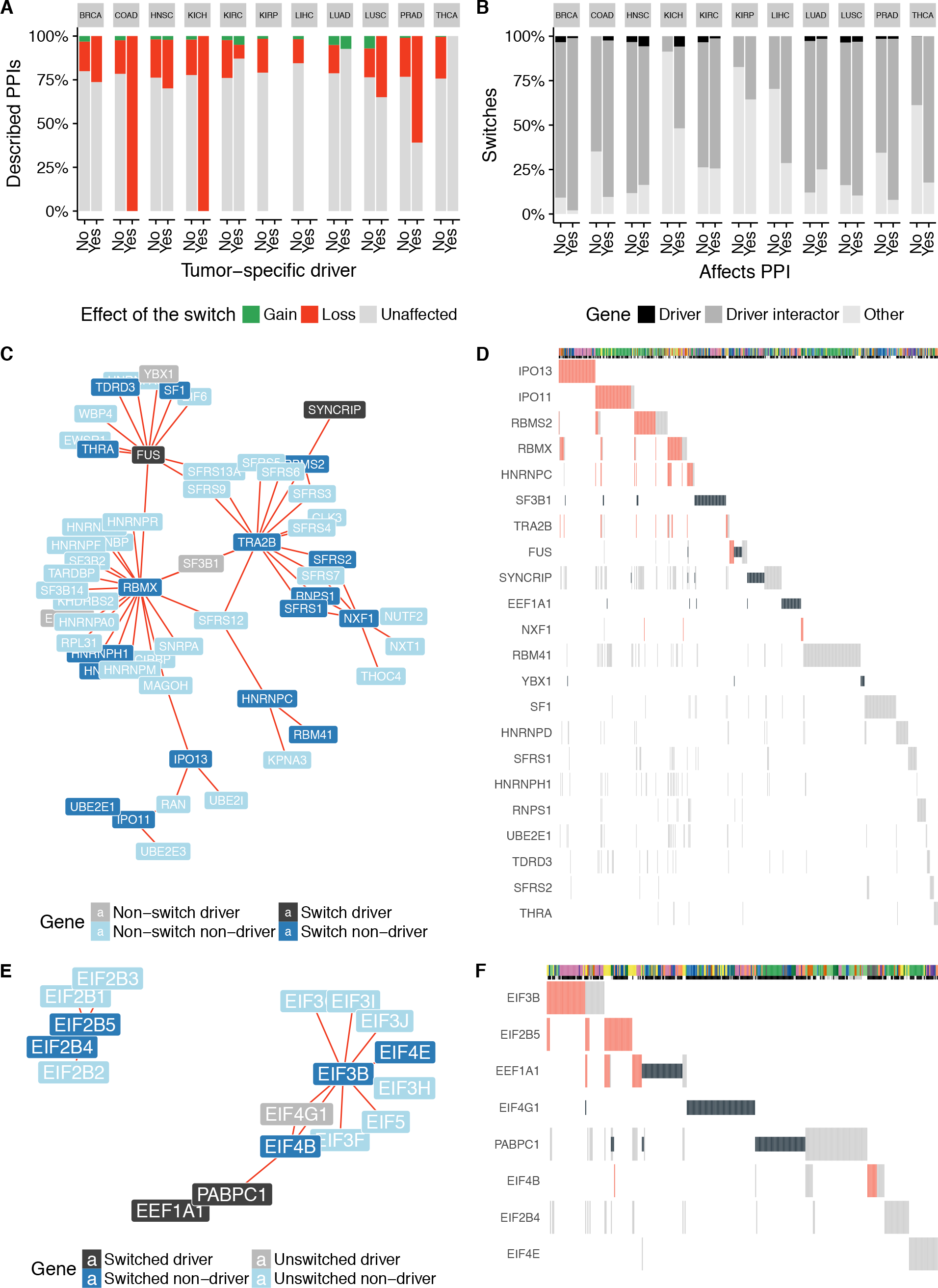
Potential impact of isoform switches in protein interactions with cancer drivers. **(A)** Functional switches were divided according to whether they occurred in tumor-specific drivers (yes) or not (no). For each tumor type we plot the proportion of protein–protein interactions (PPIs) (*y* axis) that were gained (green), lost (red), or remained unaffected (gray). All comparisons except for KIRC, LUAD were significant (Supplemental Experimental Procedures), Samples from KIRP and LIHC had no PPI-affecting switches in drivers. **(B)** Functional switches mapped to PPIs were divided according to whether they affected a PPI (yes) or not (no). For each tumor type we plotted the proportion of functional switches (*y* axis) that occurred in cancer drivers (black), in interactors of drivers (dark gray), or in other genes (light gray). All tests for the enrichment of PPIs affected by switches in driver interactors were significant except for KIRC, LUAD and LUSC (Supplemental Experimental Procedures). **(C)** Network for module 11 (Table S6) with PPIs predicted to be lost (red). Cancer drivers are indicated in black or gray if they had a functional switch or not, respectively. Other genes are indicated in dark blue or light blue if they had a functional switch or not, respectively. We do not show unaffected interactions. **(D)** OncoPrint for the samples that present protein-affecting mutations (PAMs) in drivers or switches from (C). Mutations are indicated in black and PPI-affecting switches are indicated in red (loss in this case). Other switches with no predicted effect on the PPI are depicted in gray. The top panel indicates the tumor type of each sample by color (same color code as in previous figures). The second top panel indicates whether the sample harbors a PAM in a tumor-specific driver (black) or not (gray), or whether no mutation data is available for that sample (white). **(E)** As in (C) for module 28 (Table S6). **(F)** OncoPrint for the switches and drivers from (E). Colors are as in (D).

### Isoform switches remodel protein interaction networks in cancer

To further characterize the role of switches, we calculated modules in the PPI network (Blondel et al., 2008) using only interaction edges affected by switches (Supplemental Experimental Procedures). This produced 179 modules involving 1405 genes (Table S6). From these, 52 modules included a cancer driver, and 47 of them included also switches that involved two protein-coding isoforms. We tested for the enrichment of genes belonging to specific protein complexes (Ruepp et al., 2009), complexes related to RNA-processing and splicing (Akerman et al., 2015) and cancer-related pathways (Liberzon et al., 2015) (Table S6) (Supplemental Experimental Procedures). From the 47 modules described above, 8 showed enrichment in pathways and complexes: apoptosis-related pathways (module 109 in Table S6), Ubiquitin mediated proteolysis pathway (module 26), and *ERBB* signaling pathway (module 169), as well as Spliceosomal (module 11), Ribosomal (module 170), SMN (module 28), PA700 (module 58) and TFIID (module 66) complexes (Table S6). In particular, module 11 was enriched in splicing factors and RNA binding proteins, and included the cancer drivers *SF3B1*, *FUS*, *SYNCRIP*, *EEF1A1* and *YBX1* (Figure 3C) (Table S6). The module contained a switch in *RBMX* involving the skipping of two exons and the elimination of an RNA recognition motif (RRM) that would impact interactions with *SF3B1*, *EEF1A1* and multiple RBP genes (Figure 3C); and a switch in *TRA2B* that yielded a non-coding transcript previously described (Stoilov et al., 2004) and would eliminate an interaction with *SF3B1* and other splicing factors. We also found a switch in *HNRNPC, TRA2A, NXF1 and RBMS2* that lost interactions with various SR-protein coding genes. Consistent with a potential functional impact, the PPI-affecting switches showed mutual exclusion with the mutational cancer drivers (Figure 3D). Interestingly, this module also contained switches in the Importin genes *IPO11* and *IPO13*, which affected interactions with ubiquitin conjugating enzymes *UBE2E1*, *UBE2E3* and *UBE2I*, and which showed mutual exclusion across different tumor types (Figure 3D). These results indicate that the activity of RNA-processing factors may be altered in cancer through the disruption of their PPIs by alternative splicing.

Another interesting case was module 28 (Table S6), with switches in the regulators of translation, *EIF4B*, *EIF3B* and *EIF4E*, which affected interactions with the drivers *EIF4G1*, *EIF4A2* and *PABPC1* (Figure 3E). The switch in *EIF4B* caused the skipping of one exon, which we predicted to eliminate an RRM domain and lose interactions with drivers *EIF4G1* and *PABPC1*. The switch in *EIF3B* yielded a non-coding transcript that would lose multiple interactions. Although we did not predict any PPI change for *EIF4E*, this switch lost eight predicted ANCHOR regions (Table S4), suggesting a possible effect on yet to be described interactions. Besides frequent PAMs, *PABPC1* also presented a functional switch that affected 2 disordered regions but did not affect any of the RRMs. In this case we did not predict any change in PPI and the possible functional impact remains to be discovered. Moreover, the identified PPI-affecting switches showed mutual exclusion with PAMs in *EIF4G1* and *PABPC1* and (Figure 3F). These results suggest that isoform switches may impact translational regulation in tumors through the alteration of protein–protein interactions of the corresponding regulators.

### Isoform switches as potential drivers of cancer

Our results provide evidence that a subset of the alternative splicing switches (I) induced a gain or loss of a protein domain from a family frequently mutated in cancer, (II) affected one or more PPIs, (III) displayed some mutual exclusion with drivers, or (IV) displayed recurrence across patients. One or more of these properties were fulfilled by 1662 functional switches, which we hypothesized could define potential alternative splicing drivers (potential AS-drivers) (Figure 4A) (Table S1), with the majority of them 1080 (65%) affecting mutated domain families and/or PPIs (Figure 4B). To test possible driver-like properties in these switches, we calculated their centrality and distance to mutational drivers in the PPI network, which are considered as defining properties for cancer-relevant genes (Jonsson and Bates, 2006). Potential AS-drivers showed greater centrality (Mann-Whitney test p-value < 2.2e-16) (Figure S6A) and closer distances to tumor-specific drivers (Fisher’s exact test p-value < 2.2e-16, OR = 1.5) (Figure S6B) compared to the rest of switches.

**Figure 4.**
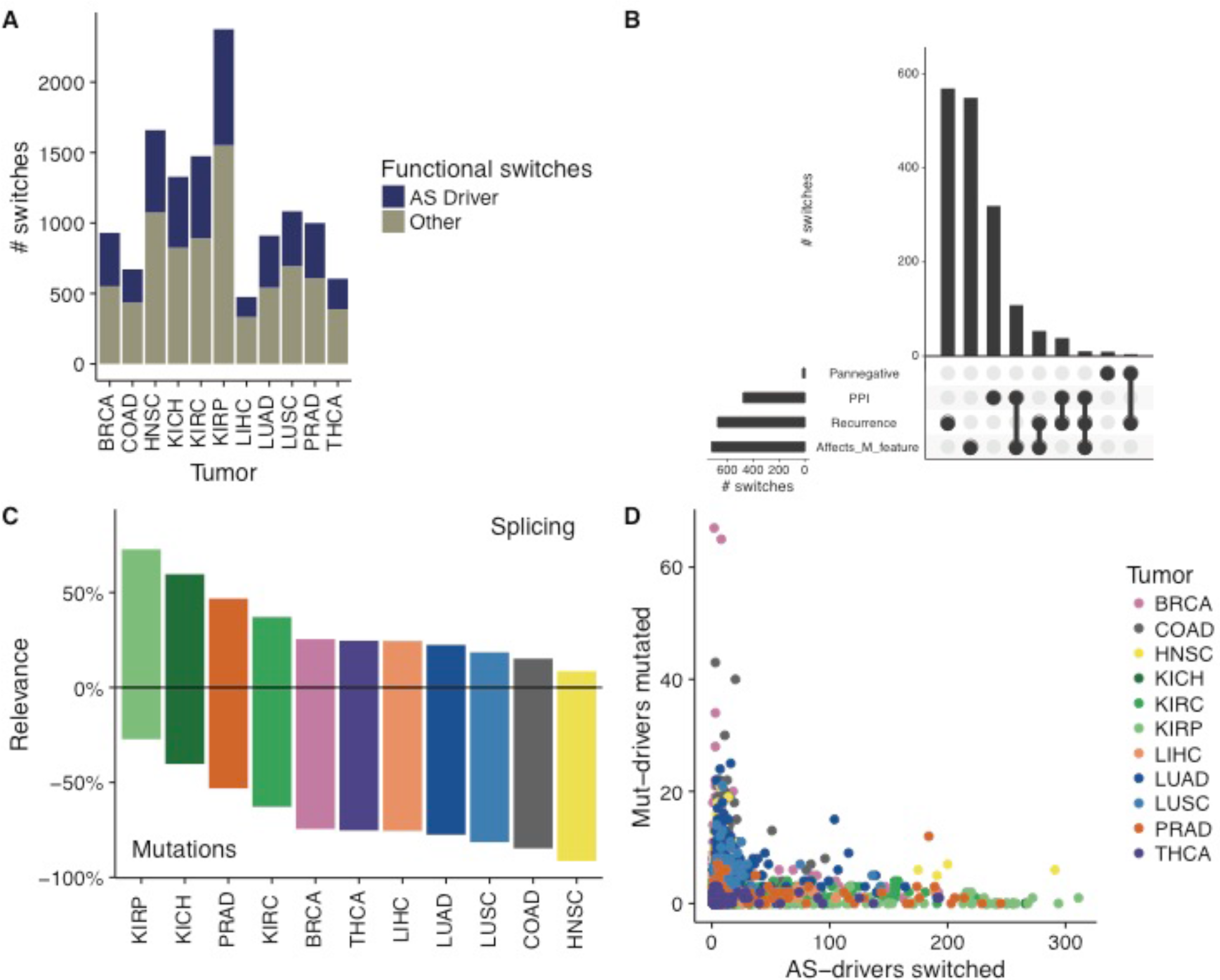
Isoform switches as potential drivers of cancer. (A) Number of functional isoform switches and potential AS-drivers detected in each tumor type. (B) Candidate potential AS-drivers grouped according to their properties: disruption of protein–protein interactions (PPIs), significant recurrence across patients (Recurrence), gain or loss of a protein feature that is frequently mutated in tumors (Affects M_feature), mutual exclusion and sharing pathway with cancer drivers (Pannegative). Horizontal bars indicate the number of switches for each property. The vertical bars indicate those in each of the intersections indicated by connected bullet points (Conway et al., 2017). (C) Classification of samples according to the relevance of potential AS-drivers or Mut-drivers in each tumor type. For each tumor type (*x* axis), the positive *y* axis shows the percentage of samples that have a proportion of switched potential AS-drivers higher than the proportion of mutated Mut-drivers. The negative *y* axis shows the percentage of samples in which the proportion of mutated Mut-drivers is higher than the proportion of switched potential AS-drivers. Only patients with mutation and transcriptome data are shown. (D) Each of the patients from (C) is represented according to the percentage of mutated Mut-drivers (*y* axis) and the percentage of switched potential AS-drivers (*x* axis).

The prevalence of these potential AS-drivers varied across samples and tumor types. Considering tumor specific mutational drivers (Mut-drivers) and our set of potential AS-drivers, we labeled each patient as AS-driver–enriched or Mut-driver–enriched according to whether the proportion of switched potential AS-drivers or mutated Mut-drivers was higher, respectively. This partition of the samples indicated that, although Mut-drivers were predominant in patients for most tumor types, potential AS-drivers were predominant for a considerable number of patients across several tumor types, and particularly for kidney and prostate tumors (Figure 4C). Additionally, regardless of the tumor type, patients with many mutations in Mut-drivers tended to show a low number of switched potential AS-drivers, and vice versa (Figure 4D). The occurrence of copy number alteration (CNA) drivers also showed a pattern of anti-correlation with our potential AS-drivers similar to the one we found between Mut-drivers and potential AS-drivers (Figure S6C). The patient distribution patterns of candidate AS-drivers compared with mutational or CNA drivers bear resemblance with the proposed cancer genome hyperbola between mutations and CNAs (Figure S6D) (Ciriello et al., 2013), which supports the notion that a subset of isoform changes represents alternative, yet-unexplored relevant mechanisms that could provide a complementary route to induce similar effects as genetic mutations.

## Discussion

We have identified consistent and recurrent transcript isoform switches that impact the function of affected proteins by adding or removing protein domains that were frequently mutated in cancer or by disrupting or gaining protein–protein interactions – possibly also altering the formation of protein complexes – with cancer drivers or in cancer related pathways. Moreover, we observed that patients with some of these isoform switches tended not to harbor mutations in cancer drivers and the other way around. Recently, an alternative splicing change in *NFE2L2* has been described to lead to the loss of a protein domain and the interaction with its negative regulator *KEAP1*, thereby providing an alternative mechanism for the activation of an oncogenic pathway (Goldstein et al., 2016). Similarly, an isoform change in the gene *ATF2* has been shown to drive melanomagenesis (Claps et al., 2016). These examples, together with the analyses presented here, support a model by which functions and pathways often altered in cancer through somatic mutations may be affected in a similar way by isoform changes in some patients, and therefore contribute to the tumor phenotype. Importantly, these isoform changes could occur without gene expression changes in the host gene and thus provide an independent catalogue of functional alterations in cancer.

Functional domains and interactions might not always be entirely lost through a switch, as normal isoforms generally retain some expression in tumors. This could be partly due to the uncertainty in the estimate of transcript abundance from RNA sequencing or to the heterogeneity in the transcriptomes of tumor cells. Still, a relatively small change in transcript abundance has ben shown to be sufficient to trigger an oncogenic effect in cells (Anczuków et al., 2015; Bechara et al., 2013; Sebestyén et al., 2016). Additionally, we observed that a number of isoform changes defined a switch from a protein-coding transcript to a non-coding one, possibly undergoing non-sense mediated decay, which is a widespread mechanism of alternative splicing mediated gene expression regulation (Hansen et al., 2009), and could potentially alter function in a way similar to other isoform changes between protein-coding isoforms. The predicted impact on domains and interactions could therefore be indicative of alterations on regulatory networks with variable functional effects.

Our description in terms of transcript isoform switches allowed us to describe more variations in the transcriptome than using local alternative splicing events, and to determine the protein features potentially gained or lost through splicing changes. However, this approach has some potential limitations. Accurate determination of differential transcript usage in genes with many isoforms requires high coverage and sufficient samples per condition (Sebestyén et al., 2015), which we expect was mitigated by our use of the variability across normal samples to determine significance. Additionally, since we used annotated transcript isoforms, we may have missed tumor specific transcripts not present in the annotation. We also only recovered a small fraction of the entire set of protein–protein interactions taking place in the cell. For instance, we did not characterize those interactions mediated through low complexity regions (Buljan et al., 2012; Ellis et al., 2012), hence many more interactions and protein complexes may be affected in tumors.

The origin of the observed splicing changes remains to be elucidated. We did not find a general association with somatic mutations in *cis*. It is possible that small copy-number alterations or *indels* are responsible for these switches, but are still hard to detect with WES and WGS data and more targeted searches or deeper sequencing is necessary. An alternative explanation is that the majority of the switches described occur through *trans* acting alterations, such as the expression change in splicing factors (Sebestyén et al., 2016). For instance, mutations in *RBM10* or downregulation of *QKI* lead to the same splicing change in *NUMB* that promotes cell proliferation (Bechara et al., 2013; Zong et al., 2014), and the oncogenic switch in *RAC1* (Zhou et al., 2012) is regulated by expression changes in various splicing factors (Gonçalves et al., 2009; Pelisch et al., 2012), which are controlled by pathways often altered in tumors (Fu and Ares, 2014). Another possibility is that these switches describe signatures of non-genetic variability (Brock et al., 2009). The intra-tumor heterogeneity could allow recapitulating similar transcriptome phenotypes, which would determine the fitness of cells and the progression of tumors independently of somatic mutations. Since natural selection acts on the phenotype rather than on the genotype, an interesting hypothesis is that specific transcript isoform expression patterns could define particular tumor phenotypes that would be closely related to those determined by somatic mutations in drivers, thereby defining an advantageous phenotype such that the selective pressure to develop equivalent adaptations is relaxed. Accordingly, our identified isoform switches could play an important role in the neoplastic process independently of or in conjunction with the already characterized genetic alterations.

## Experimental Procedures

Further details and an outline of resources used in this work can be found in Supplemental Experimental Procedures.

### Calculation of significant isoform switches per patient

We modeled splicing alterations in a gene as a switch between two transcript isoforms, one normal and one tumoral. For each transcript, the relative abundance per sample, which we called PSI, was calculated by normalizing its abundance in TPM units by the sum of abundances of all transcripts in the same gene. Then, for each transcript and sample we calculated the change in relative abundance as ΔPSI = PSI_tumor_-PSI_ref_, where PSI_tumor_ is the relative abundance in the tumor sample and PSIref is the normal reference value, which is the value of the paired normal sample when available, or the median of PSIs in the normal samples for the same tissue type, otherwise. We considered significant those changes with |ΔPSI| > 0.05 and with empirical p < 0.01 in the comparison of the observed |ΔPSI| value with the distribution of |ΔPSI| values obtained by comparing the normal samples pairwise without repetition. We only kept those cases for which the tumor isoform PSI was higher than the normal isoform in the tumor sample and the normal isoform PSI in the normal sample was higher than the value for the tumor isoform. Moreover, we discarded genes that either had an outlier expression in the tumor sample compared to normal tissues – had expression below the bottom 2.5% or above the 97.5% of the values of normal expression – or showed differential expression between the tumor and the normal samples (Wilcoxon test p-value < 0.01 using the gene TPM values).

Candidate switches were defined per patient and per gene, and in some samples the same gene could have different switches. We discarded those switches that contradicted a more frequent switch in the same gene and the same tumor type. Moreover, we discarded any switch that affected a number of patients below the top 99% of the distribution of patient frequency of these contradictory switches in each tumor type. Lastly, we filtered out switches that were significantly lowly recurrent, i.e. they occurred in fewer patients than expected by chance (Binomial test - adjusted p-value < 0.05, using all tumor types). As a consequence, none of the reported switches occurred in less than 5 samples. Thus, a switch in a patient sample was defined as a pair of transcripts in a gene with no expression change and with significant changes in opposite directions that showed consistency across a minimum number of patients. We aggregated the switches from the different tumor types to get the final list (Table S1).

### Simulated switches

To simulate switches between normal and tumor tissues we used genes with more than one expressed isoform. For each gene, we selected the isoform with the highest median expression across the normal samples as the normal isoform and an arbitrary different transcript expressed in the tumor samples as the tumor isoform. For each gene, we generated a maximum of five such simulated switches.

### Functional switches

A switch was defined as functional if both isoforms overlapped in genomic extent and there was a change in the encoded protein, including cases where only one of the isoforms was coding, and moreover there was a gain or loss of a protein feature: Pfam domains (Finn et al., 2016) mapped with InterProScan (Jones et al., 2014), ProSite patterns (Gattiker et al., 2002); disordered regions from IUPred (Dosztanyi et al., 2005); and disordered regions potentially involved in protein–protein interactions from ANCHOR (Dosztanyi et al., 2009). For IUPred and ANCHOR we only considered changes involving at least 5 amino acids. Switches without any mapped protein features were not considered. Significance on the enrichment of protein features losses versus gains was calculated by comparing the number of gains and losses in switches with the same numbers in simulated switches (Supplemental Experimental Procedures).

### Enrichment of domain families in switches and mutations

To find protein domain families significantly affected by switches we first calculated a reference proteome for each tumor type. Using genes with multiple transcripts, we selected those that had at least one isoform with TPM>0.1, and only kept the isoform with the highest median expression across the normal samples in the same tissue type. Proteins encoded by these isoforms were considered the reference proteome in each tumor type. We aggregated the reference proteomes from all tumor types to form a pan-cancer reference proteome. The expected frequency of a protein feature was then measured as the proportion of this feature in the reference proteome. This expected frequency was then used to calculate the probability of a feature to be affected by a switch using a binomial test with the number of times the feature was gained or lost in switches and the total number of feature gains or losses due to switches (Supplemental Experimental Procedures). We selected cases with Benjamini-Hochberg (BH) adjusted p-value < 0.05. Additionally, to ensure the specificity of the enrichment for each domain class, we considered only domain families affected in at least two switches. To calculate domain families enriched in mutations, we considered again the reference proteome in each tumor type. The expected mutation rate of a domain family was considered to be the proportion of the length of domains in the proteome covered by this domain family. We aggregated all observed mutations falling within each family and calculated the probability of the observed mutations using a binomial test using the mutation count for a domain family and the total mutations in all domain families (Supplemental Experimental Procedure). After correcting for multiple testing, we kept those cases with a BH adjusted p-value < 0.05. GO analysis was performed using DcGO (Fang and Gough, 2013). For the enrichment test, we considered significant those cases with FDR < 0.01 (hypergeometric test).

### Protein interaction analysis

We created a consensus protein–protein interaction (PPI) network using data from PSICQUIC (del Toro et al. 2013), BIOGRID (Chatr-Aryamontri et al., 2015), HumNet (Lee et al., 2011), STRING (Szklarczyk et al., 2011), and from (Rolland et al., 2014). The consensus network was built with interactions appearing in at least four of these five sources, yielding a total of 8,142 nodes with 29,991 interactions. To find PPIs likely altered by isoform switches we first mapped each PPI in a gene to a specific domain–domain interaction (DDI), using information on domain–domain interactions from iPfam (Finn et al., 2014), DOMINE (Raghavachari et al., 2008), and 3did (Mosca et al., 2014). Domains involved in DDIs were then mapped to specific protein isoforms. For the genes with switches, we then considered those PPIs that could be mapped to DDIs involving domains mapped to either the normal or the tumor isoforms. In total, 3,242 genes with 4,219 switches mapped to one or more interactions in the consensus network, and 1,688 isoform switches (in 1,355 genes) were mapped to at least one specific DDI. We defined a PPI as lost if it was mapped to one or more DDIs in the isoform expressed in the normal tissue but not in the isoform expressed in the tumor sample. If multiple domains mediated the same interaction, it was considered lost if at least one of these domains was lost in the switch. We defined a PPI as gained if it was mapped to a DDI only in the tumor isoform but not in the normal isoform.

## Author contributions

E.E. proposed the study. H.C-G developed the software and performed the analyses. E.P-P built the consensus protein–protein interaction network and mapped the domain–domain interactions. E.E. and A.G. supervised the analyses. E.E. and H.C-G wrote the manuscript with essential inputs from E.P-P and A.G.

## Acknowledgements

H.C-G and E.E. were supported by the MINECO and FEDER (BIO2014-52566-R), Consolider RNAREG (CSD2009-00080), AGAUR (SGR2014-1121), the European ITN Network RNP-Net (ID:289007) and the Sandra Ibarra Foundation for Cancer (FSI2013). E.P-P and A.G. were supported by the SBP CC grant (P30 CA030199). All authors thank The Cancer Genome Atlas project for making their data publicly available.

## References

Akerman, M., Fregoso, O.I., Das, S., Ruse, C., Jensen, M. a, Pappin, D.J., Zhang, M.Q., and Krainer, A.R. (2015). Differential connectivity of splicing activators and repressors to the human spliceosome. Genome Biol. 16, 119.

Alamancos, G.P., Pagés, A., Trincado, J.L., Bellora, N., and Eyras, E. (2015). Leveraging transcript quantification for fast computation of alternative splicing profiles. RNA 21, 1521–1531.

Alsafadi, S., Houy, A., Battistella, A., Popova, T., Wassef, M., Henry, E., Tirode, F., Constantinou, A., Piperno-Neumann, S., Roman-Roman, S., et al. (2016). Cancer-associated SF3B1 mutations affect alternative splicing by promoting alternative branchpoint usage. Nat. Commun. 7, 10615.

Anczuków, O., Akerman, M., Cléry, A., Wu, J., Shen, C., Shirole, N.H., Raimer, A., Sun, S., Jensen, M.A., Hua, Y., et al. (2015). SRSF1-Regulated Alternative Splicing in Breast Cancer. Mol. Cell 60, 105–117.

Bechara, E.G., Sebestyén, E., Bernardis, I., Eyras, E., and Valcárcel, J. (2013). RBM5, 6, and 10 differentially regulate NUMB alternative splicing to control cancer cell proliferation. Mol. Cell 52.

Birzele, F., Csaba, G., and Zimmer, R. (2008). Alternative splicing and protein structure evolution. Nucleic Acids Res. 36, 550–558.

Blondel, V.D., Guillaume, J.-L., Lambiotte, R., and Lefebvre, E. (2008). Fast unfolding of communities in large networks. J. Stat. Mech. Theory Exp. 10008, 6.

Bourdon, J.-C. (2007). P53 and Its Isoforms in Cancer. Br. J. Cancer 97, 277–282.

Brock, A., Chang, H., and Huang, S. (2009). Non-genetic heterogeneity—a mutation-independent driving force for the somatic evolution of tumours. Nat. Rev. Genet. 10, 336–342.

Buljan, M., Chalancon, G., Eustermann, S., Wagner, G.P., Fuxreiter, M., Bateman, A., and Babu, M.M. (2012). Tissue-Specific Splicing of Disordered Segments that Embed Binding Motifs Rewires Protein Interaction Networks. Mol. Cell 46, 871–883.

Chabot, B., and Shkreta, L. (2016). Defective control of pre-messenger RNA splicing in human disease. J. Cell Biol. 212, 13–27.

Chatr-Aryamontri, A., Breitkreutz, B.J., Oughtred, R., Boucher, L., Heinicke, S., Chen, D., Stark, C., Breitkreutz, A., Kolas, N., O’Donnell, L., et al. (2015). The BioGRID interaction database: 2015 update. Nucleic Acids Res. 43, D470–D478.

Ciriello, G., Miller, M.L., Aksoy, B.A., Senbabaoglu, Y., Schultz, N., and Sander, C. (2013). Emerging landscape of oncogenic signatures across human cancers. Nat. Genet. 45, 1127–1133.

Claps, G., Cheli, Y., Zhang, T., Scortegagna, M., Lau, E., Kim, H., Qi, J., Li, J.-L., James, B., Dzung, A., et al. (2016). A Transcriptionally Inactive ATF2 Variant Drives Melanomagenesis. Cell Rep. 15, 1884–1892.

Conway, J.R., Lex, A., and Gehlenborg, N. (2017). UpSetR: An R Package for the Visualization of Intersecting Sets and their Properties. 2–5.

Darman, R.B., Seiler, M., Agrawal, A.A., Lim, K.H., Peng, S., Aird, D., Bailey, S.L., Bhavsar, E.B., Chan, B., Colla, S., et al. (2015). Cancer-Associated SF3B1 Hotspot Mutations Induce Cryptic 3′ Splice Site Selection through Use of a Different Branch Point. Cell Rep. 13, 1033–1045.

David, C.J., and Manley, J.L. (2010). Alternative pre-mRNA splicing regulation in cancer: Pathways and programs unhinged. Genes Dev. 24, 2343–2364.

Dosztányi, Z., Csizmok, V., Tompa, P., and Simon, I. (2005). IUPred: web server for the prediction of intrinsically unstructured regions of proteins based on estimated energy content. Bioinformatics 21, 3433–3434.

Dosztányi, Z., Mészáros, B., and Simon, I. (2009). ANCHOR: web server for predicting protein binding regions in disordered proteins. Bioinformatics 25, 2745–2746.

Dvinge, H., and Bradley, R.K. (2015). Widespread intron retention diversifies most cancer transcriptomes. Genome Med. 7, 45.

Ellis, J.D., Barrios-Rodiles, M., ??olak, R., Irimia, M., Kim, T., Calarco, J.A., Wang, X., Pan, Q., O’Hanlon, D., Kim, P.M., et al. (2012). Tissue-Specific Alternative Splicing Remodels Protein-Protein Interaction Networks. Mol. Cell 46, 884–892.

Fang, H., and Gough, J. (2013). DcGO: Database of domain-centric ontologies on functions, phenotypes, diseases and more. Nucleic Acids Res. 41.

Finn, R.D., Miller, B.L., Clements, J., and Bateman, A. (2014). IPfam: A database of protein family and domain interactions found in the Protein Data Bank. Nucleic Acids Res. 42.

Finn, R.D., Coggill, P., Eberhardt, R.Y., Eddy, S.R., Mistry, J., Mitchell, A.L., Potter, S.C., Punta, M., Qureshi, M., Sangrador-Vegas, A., et al. (2016). The Pfam protein families database: towards a more sustainable future. Nucleic Acids Res. 44, D279–85.

Frampton, G.M., Ali, S.M., Rosenzweig, M., Chmielecki, J., Lu, X., Bauer, T.M., Akimov, M., Bufill, J.A., Lee, C., Jentz, D., et al. (2015). Activation of MET via diverse exon 14 splicing alterations occurs in multiple tumor types and confers clinical sensitivity to MET inhibitors. Cancer Discov. 5, 850–860.

Fu, X.-D., and Ares, M. (2014). Context-dependent control of alternative splicing by RNA-binding proteins. Nat. Rev. Genet. 15, 689–701.

Gattiker, A., Gasteiger, E., and Bairoch, A. (2002). ScanProsite: a reference implementation of a PROSITE scanning tool. Appl. Bioinformatics 1, 107–108.

Goldstein, L.D., Lee, J., Gnad, F., Klijn, C., Schaub, A., Reeder, J., Daemen, A., Bakalarski, C.E., Holcomb, T., Shames, D.S., et al. (2016). Recurrent Loss of NFE2L2 Exon 2 Is a Mechanism for Nrf2 Pathway Activation in Human Cancers. Cell Rep. 16, 2605–2617.

Gonçalves, V., Matos, P., and Jordan, P. (2009). Antagonistic SR proteins regulate alternative splicing of tumor-related Rac1b downstream of the PI3-kinase and Wnt pathways. Hum. Mol. Genet. 18, 3696–3707.

Hansen, K.D., Lareau, L.F., Blanchette, M., Green, R.E., Meng, Q., Rehwinkel, J., Gallusser, F.L., Izaurralde, E., Rio, D.C., Dudoit, S., et al. (2009). Genome-wide identification of alternative splice forms down-regulated by nonsense-mediated mRNA decay in Drosophila. PLoS Genet. 5.

Jones, P., Binns, D., Chang, H.Y., Fraser, M., Li, W., McAnulla, C., McWilliam, H., Maslen, J., Mitchell, A., Nuka, G., et al. (2014). InterProScan 5: Genome-scale protein function classification. Bioinformatics 30, 1236–1240.

Jonsson, P.F., and Bates, P.A. (2006). Global topological features of cancer proteins in the human interactome. Bioinformatics 22, 2291–2297.

Jung, H., Lee, D., Lee, J., Park, D., Kim, Y.J., Park, W.-Y., Hong, D., Park, P.J., and Lee, E. (2015). Intron retention is a widespread mechanism of tumor-suppressor inactivation. Nat. Genet. 47, 1242–1248.

Karni, R., de Stanchina, E., Lowe, S.W., Sinha, R., Mu, D., and Krainer, A.R. (2007). The gene encoding the splicing factor SF2/ASF is a proto-oncogene. Nat. Struct. Mol. Biol. 14, 185–193.

Lee, S.C.-W., and Abdel-Wahab, O. (2016). Therapeutic targeting of splicing in cancer. Nat. Med. 22, 976–986.

Lee, I., Blom, U.M., Wang, P.I., Shim, J.E., and Marcotte, E.M. (2011). Prioritizing candidate disease genes by network-based boosting of genome-wide association data. Genome Res. 21, 1109–1121.

Liberzon, A., Birger, C., Thorvaldsdóttir, H., Ghandi, M., Mesirov, J.P., and Tamayo, P. (2015). The Molecular Signatures Database Hallmark Gene Set Collection. Cell Syst. 1, 417–425.

Lu, Z., Huang, Q., Park, J.W., Shen, S., Lin, L., Tokheim, C.J., Henry, M.D., and Xing, Y. (2015). Transcriptome-wide landscape of pre-mRNA alternative splicing associated with metastatic colonization. Mol. Cancer Res. 13, 305–318.

Madan, V., Kanojia, D., Li, J., Okamoto, R., Sato-Otsubo, A., Kohlmann, A., Sanada, M., Grossmann, V., Sundaresan, J., Shiraishi, Y., et al. (2015). Aberrant splicing of U12-type introns is the hallmark of ZRSR2 mutant myelodysplastic syndrome. Nat. Commun. 6, 6042.

Mosca, R., Céol, A., Stein, A., Olivella, R., and Aloy, P. (2014). 3did: A catalog of domain-based interactions of known three-dimensional structure. Nucleic Acids Res. 42.

Norris, A.D., and Calarco, J.A. (2012). Emerging roles of alternative pre-mRNA splicing regulation in neuronal development and function. Front. Neurosci.

Paik, P.K., Drilon, A., Fan, P.-D., Yu, H., Rekhtman, N., Ginsberg, M.S., Borsu, L., Schultz, N., Berger, M.F., Rudin, C.M., et al. (2015). Response to MET Inhibitors in Patients with Stage IV Lung Adenocarcinomas Harboring MET Mutations Causing Exon 14 Skipping. Cancer Discov. 5, 842–849.

Pelisch, F., Khauv, D., Risso, G., Stallings-Mann, M., Blaustein, M., Quadrana, L., Radisky, D.C., and Srebrow, A. (2012). Involvement of hnRNP A1 in the matrix metalloprotease-3-dependent regulation of Rac1 pre-mRNA splicing. J. Cell. Biochem. 113, 2319–2329.

Poulikakos, P.I., Persaud, Y., Janakiraman, M., Kong, X., Ng, C., Moriceau, G., Shi, H., Atefi, M., Titz, B., Gabay, M.T., et al. (2011). RAF inhibitor resistance is mediated by dimerization of aberrantly spliced BRAF(V600E). Nature 480, 387–390.

Raghavachari, B., Tasneem, A., Przytycka, T.M., and Jothi, R. (2008). DOMINE: A database of protein domain interactions. Nucleic Acids Res. 36.

Rolland, T., Taşan, M., Charloteaux, B., Pevzner, S.J., Zhong, Q., Sahni, N., Yi, S., Lemmens, I., Fontanillo, C., Mosca, R., et al. (2014). A proteome-scale map of the human interactome network. Cell 159, 1212–1226.

Ruepp, A., Waegele, B., Lechner, M., Brauner, B., Dunger-Kaltenbach, I., Fobo, G., Frishman, G., Montrone, C., and Mewes, H.W. (2009). CORUM: The comprehensive resource of mammalian protein complexes-2009. Nucleic Acids Res. 38.

Saito, M., Shimada, Y., Shiraishi, K., Sakamoto, H., Tsuta, K., Totsuka, H., Chiku, S., Ichikawa, H., Kato, M., Watanabe, S.I., et al. (2015). Development of lung adenocarcinomas with exclusive dependence on oncogene fusions. Cancer Res. 75, 2264–2271.

Sebestyén, E., Zawisza, M., and Eyras, E. (2015). Detection of recurrent alternative splicing switches in tumor samples reveals novel signatures of cancer. Nucleic Acids Res. 43, 1345–1356.

Sebestyén, E., Singh, B., Miñana, B., Pagès, A., Mateo, F., Pujana, M.A., Valcárcel, J., and Eyras, E. (2016). Large-scale analysis of genome and transcriptome alterations in multiple tumors unveils novel cancer-relevant splicing networks. Genome Res. 26, 732–744.

Sotillo, E., Barrett, D.M., Black, K.L., Bagashev, A., Oldridge, D., Wu, G., Sussman, R., Lanauze, C., Ruella, M., Gazzara, M.R., et al. (2015). Convergence of acquired mutations and alternative splicing of CD19 enables resistance to CART-19 immunotherapy. Cancer Discov. 5, 1282–1295.

Stoilov, P., Daoud, R., Nayler, O., and Stamm, S. (2004). Human tra2-beta1 autoregulates its protein concentration by influencing alternative splicing of its pre-mRNA. Hum. Mol. Genet. 13, 509–524.

Supek, F., Miñana, B., Valcárcel, J., Gabaldón, T., and Lehner, B. (2014). Synonymous mutations frequently act as driver mutations in human cancers. Cell 156, 1324–1335.

Szklarczyk, D., Franceschini, A., Kuhn, M., Simonovic, M., Roth, A., Minguez, P., Doerks, T., Stark, M., Muller, J., Bork, P., et al. (2011). The STRING database in 2011: Functional interaction networks of proteins, globally integrated and scored. Nucleic Acids Res. 39.

Trincado, J.L., Sebestyen, E., Pages, A., Eyras, E., Sebestyén, E., Pagés, A., and Eyras, E. (2016). The prognostic potential of alternative transcript isoforms across human tumors. Genome Med. 8, 85.

Vorlová, S., Rocco, G., Lefave, C. V, Jodelka, F.M., Hess, K., Hastings, M.L., Henke, E., and Cartegni, L. (2011). Induction of antagonistic soluble decoy receptor tyrosine kinases by intronic polyA activation. Mol. Cell 43, 927–939.

Wang, P., Yan, B., Guo, J.-T., Hicks, C., and Xu, Y. (2005). Structural genomics analysis of alternative splicing and application to isoform structure modeling. Proc. Natl. Acad. Sci. U. S. A. 102, 18920–18925.

Yanagisawa, M., Huveldt, D., Kreinest, P., Lohse, C.M., Cheville, J.C., Parker, A.S., Copland, J.A., and Anastasiadis, P.Z. (2008). A p120 catenin isoform switch affects rho activity, induces tumor cell invasion, and predicts metastatic disease. J. Biol. Chem. 283, 18344–18354.

Yang, F., Petsalaki, E., Rolland, T., Hill, D.E., Vidal, M., and Roth, F.P. (2015). Protein Domain-Level Landscape of Cancer-Type-Specific Somatic Mutations. PLoS Comput. Biol. 11.

Yang, X., Coulombe-Huntington, J., Kang, S., Sheynkman, G.M., Hao, T., Richardson, A., Sun, S., Yang, F., Shen, Y.A., Murray, R.R., et al. (2016). Widespread Expansion of Protein Interaction Capabilities by Alternative Splicing. Cell 164, 805–817.

Zhou, C., Licciulli, S., Avila, J.L., Cho, M., Troutman, S., Jiang, P., Kossenkov, a V, Showe, L.C., Liu, Q., Vachani, a, et al. (2012). The Rac1 splice form Rac1b promotes K-ras-induced lung tumorigenesis. Oncogene 32, 903–909.

Zong, F.Y., Fu, X., Wei, W.J., Luo, Y.G., Heiner, M., Cao, L.J., Fang, Z., Fang, R., Lu, D., Ji, H., et al. (2014). The RNA-Binding Protein QKI Suppresses Cancer-Associated Aberrant Splicing. PLoS Genet. 10.

